# A protein semisynthesis-based strategy to probe the impact of site-specific serine ADP-ribosylation on linker histone function

**DOI:** 10.1101/2022.02.17.480967

**Authors:** Kyuto Tashiro, Jugal Mohapatra, Chad A. Brautigam, Glen Liszczak

## Abstract

Recently developed chemical and enzyme-based technologies to install serine ADP-ribosylation onto synthetic peptides have enabled new approaches to study PARP biology. Here, we establish a generalizable strategy to prepare ADP-ribosylated peptides that are compatible with N-terminal, C-terminal and sequential protein ligation reactions. Two unique protein-assembly routes are employed to generate full-length linker histone constructs that are homogenously ADP-ribosylated at known DNA damage-dependent modification sites. We found that serine mono-ADP-ribosylation is sufficient to alleviate linker histone-dependent chromatin compaction, and that this effect is amplified by ADP-ribose chain elongation. Our work will greatly expand the scope of ADP-ribose-modified proteins that can be constructed via semisynthesis, which is rapidly emerging as a robust approach to elucidate the direct effects that site-specific serine mono- and poly-ADP-ribosylation have on protein function.

---

Protein ADP-ribosylation (ADPr) is an NAD^+^-dependent post-translational modification that targets hundreds of proteins in mammalian cells to regulate diverse signaling processes^1^. Among the most widespread forms of this modification is DNA damage-induced serine ADPr, which is catalyzed by the PARP1/2:HPF1 complex^2–5^. While intense efforts have been directed towards understanding PARP1/2:HPF1 regulatory mechanisms^6–8^, substrate preferences^4,9,10^, and factors that govern ADP-ribose chain elongation^11–13^ many questions remain surrounding how specific mono- and poly-ADPr events affect target protein function. A greater understanding of ADPr-mediated signaling mechanisms could expand the effective use of PARP1/2 inhibitors for the treatment of homology repair-deficient cancers and other diseases^14^, and has the potential to uncover alternative treatment strategies.

We and others recently developed synthetic and chemoenzymatic strategies to prepare peptides bearing site-specific serine ADPr^12,15,16^. These approaches are compatible with peptide thioesters to enable native chemical ligation, which is a cornerstone of protein post-translational modification research that can now be applied to the study of serine ADPr. However, the core histones H2B and H3 remain the only two full-length, serine ADP-ribosylated proteins that have been assembled via semisynthesis^12,17^. Notably, both H2B and H3 were prepared via a single ligation reaction wherein the ADP-ribosylated N-terminal peptide thioester fragment was ligated to a recombinant C-terminal fragment. More modular functionalization of ADP-ribosylated peptides, including an N-terminal cysteine and a latent C-terminal thioester, is necessary to grant access to modification sites throughout an entire protein sequence. This design would vastly expand the number of semisynthetically modified PARP1/2:HPF1 substrates that can be functionally characterized in biochemical, biophysical, and cell-based assays.

The disordered, lysine-rich C-terminal domain of linker histone H1 contains several serine residues that act as acceptor sites for DNA damage-induced ADPr^4,8,9,18^. It is clear that PARP1/2 activity is required to release linker histone H1 from chromatin at DNA damage sites, which contributes to local chromatin relaxation and DNA repair^18,19^. Multiple mechanisms to explain how PARP1 activity impacts H1 function have been put forth, including: (i) PARP1 displaces H1 from chromatin by engaging an overlapping nucleosome binding site, (ii) a non-covalent H1:poly-ADPr interaction disrupts the H1:DNA interface, and (iii) poly-anionic ADP-ribose chains reduce the affinity between the highly basic H1 C-terminus and the negatively charged chromatin polymer^19–22^. Additionally, H1 ADPr may induce downstream protein-modification cascades or ADP-ribose-binding protein recruitment events that influence H1 function. With semisynthetic ADP-ribosylated H1 constructs, chromatin structure analysis can be performed in the absence of confounding variables that include the PARP1 enzyme, core histone ADPr, and other protein modifications or DNA repair factors. Thus, the impact that site-specific mono- and poly-ADPr have on H1 function can be directly interrogated.

We chose to study the linker histone H1.2 as it is an abundant variant and a known target of DNA damage-induced serine ADPr^8,18^. There are four PARP1/2:HPF1 substrate motifs (Lys-Ser) in H1.2 (Figure 1A), of which several have been reported as ADPr acceptor sites in mammalian cell-based assays. To determine the primary H1.2 ADPr sites, we introduced wild-type or serine-to-alanine 6xHis-tagged H1.2 transgenes into HEK293T cells for ADPr analysis. Following exposure to an oxidative DNA damage agent, cells were collected and lysed in a denaturing buffer that included 6M urea to rapidly quench all ADPr and glycohydrolase activity. Lysates were then cleared and passed over an Ni-NTA column to enrich H1.2 transgenes for western blot analysis (Figure 1B). We found that all detectable H1.2 ADPr occurs at the S150 and S188 sites, as confirmed by the S150A/S188A double mutant. This modification site preference is also maintained in reconstituted ADPr assays comprising the PARP1/2:HPF1 complex and recombinant, full-length H1.2 constructs (Figure 1C). While the S86 site falls within the folded domain of H1.2 and is likely sterically protected from PARP1 activity, the S150, S173 and S188 sites are located within the C-terminal disordered region. Unlike S150 and S188, the S173 site is immediately followed by a proline residue, which we hypothesized may prevent modification by the PARP1:HPF1 complex. To test this concept, an RP-HPLC/MS-based assay was employed to analyze PARP1:HPF1 activity on H1.2 peptide fragments (Figure 1D, Figure S1). Indeed, while ADPr could not be detected on the H1.2_166-181_ fragment, the H1.2_143-158_, H1.2_182-195_, and an H1.2_166-181_ bearing a P174A mutation were quantitatively ADP-ribosylated after a 20 minute incubation with PARP1:HPF1 (1 μM:25 μM) at 30 °C. Thus, a proline residue C-terminal to the Lys-Ser motif potently inhibits PARP1:HPF1 activity.

**Figure 1.**
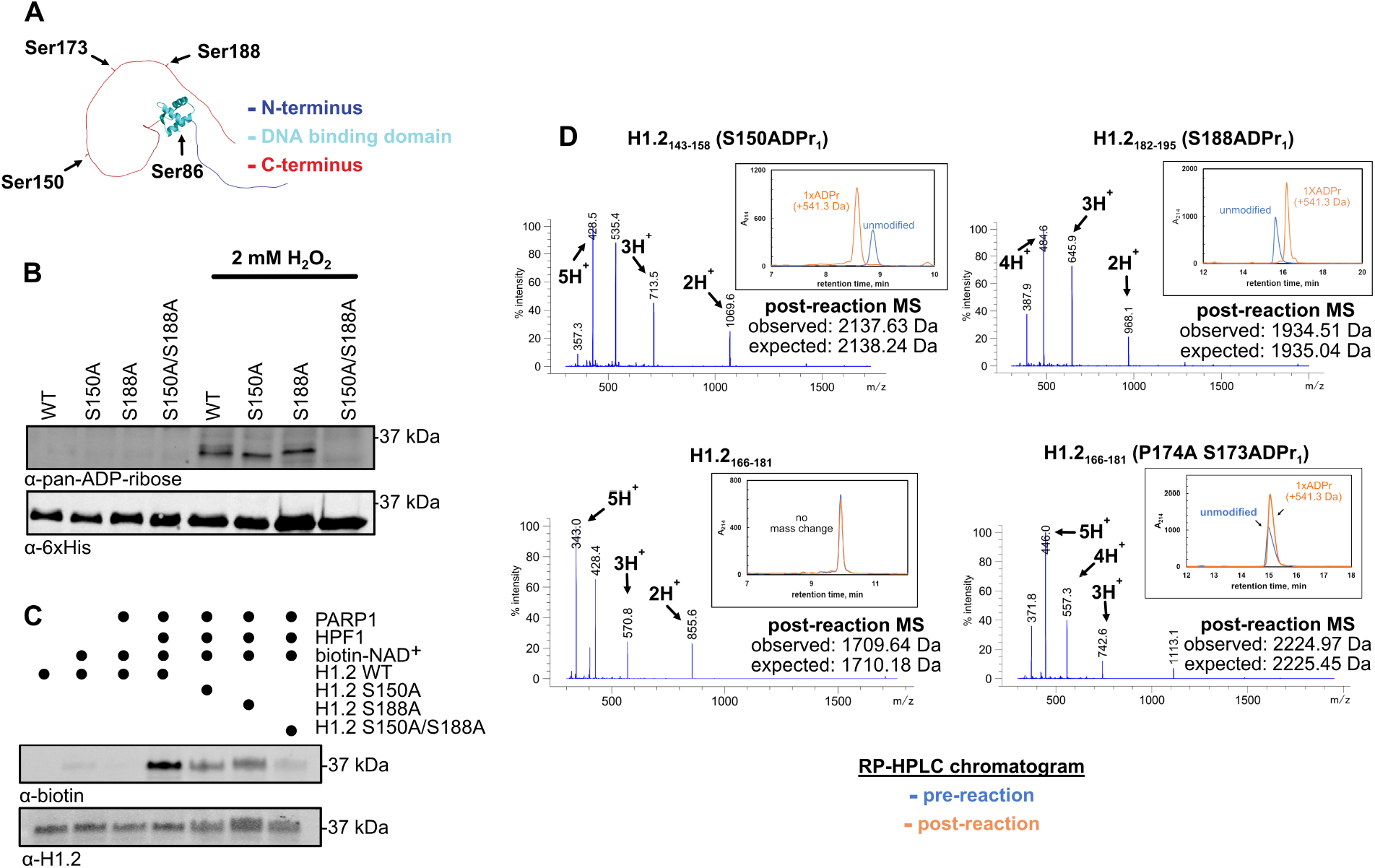
Histone H1.2 S150 and S188 are the main DNA-damage dependent ADP-ribosylation sites. (A) AlphaFold-predicted three-dimensional structure of full-length H1.2 with serine residue that occur within a Lys-Ser motif indicated. (B) Western blot analysis of the indicated 6xHis-tagged H1.2 transgenes that were enriched from HEK293T cells via Ni-NTA affinity pre- and post-exposure to hydrogen peroxide (2 mM, 15 min). (C) Western blot analysis of reconstituted, biotin-NAD^+^-based PARP1:HPF1 ADPr assays consisting of the indicated recombinant enzymatic components and full-length H1.2 constructs. (D) RP-HPLC analysis of the indicated synthetic H1.2 peptide fragments pre- and post-incubation with reconstituted PARP1:HPF1 and NAD^+^. Intact ESI-MS analysis of each post-reaction H1.2 fragment is shown.

With key H1.2 serine ADPr target sites established, we set out to prepare semisynthetic H1.2 constructs modified with mono-ADP-ribose at the S150 or S188 site. We envisioned a three-piece native chemical ligation strategy to gain synthetic access to the S150 site (Figure 2A). A recombinant piece 1 thioester fragment (amino acids 2-141; H1.2_2-141_) was prepared via an intein fusion-based approach and a recombinant piece 3 fragment (amino acids 163-213, A163C; H1.2_163-213_) was prepared via standard procedures (Figure S2). The synthetic piece 2 fragment (amino acids 142-162, A142C; H1.2_142-162_) was initially functionalized with an N-terminal cysteine and a C-terminal acyl hydrazide, which can be converted to a thioester in a biorthogonal reaction^23^ to enable sequential protein ligations starting from the N-terminus. Following peptide synthesis, H1.2_142-162_ was incubated with the PARP1:HPF1 complex in the presence of PARG to install the mono-ADP-ribose modification at S150, the sole serine residue in the peptide. However, post-reaction analysis revealed that the H1.2_142-162_ fragment is modified with two ADP-ribose moieties, and a similar peptide substrate bearing an S150A mutation maintained a single PARP1:HPF1 modification site (Figure S3). Considering that the highly similar H1.2_143-158_ S150A construct was not a PARP1:HPF1 substrate, we hypothesized that the H1.2_142-162_ secondary modification site is dependent upon the N-terminal cysteine residue. To unambiguously identify the modification site, we incubated the unmodified S150A mutant peptide bearing an N-terminal cysteine and the ADP-ribosylated variant of this peptide with iodoacetamide. Interestingly, while the unmodified S150A mutant could be labeled with iodoacetamide as confirmed by RP-HPLC/MS analysis, the ADP-ribosylated variant of this peptide was resistant to iodoacetamide treatment (Figure 2B and S3). Therefore, the PARP1:HPF1 complex modifies the thiol moiety of N-terminal cysteine residues.

**Figure 2.**
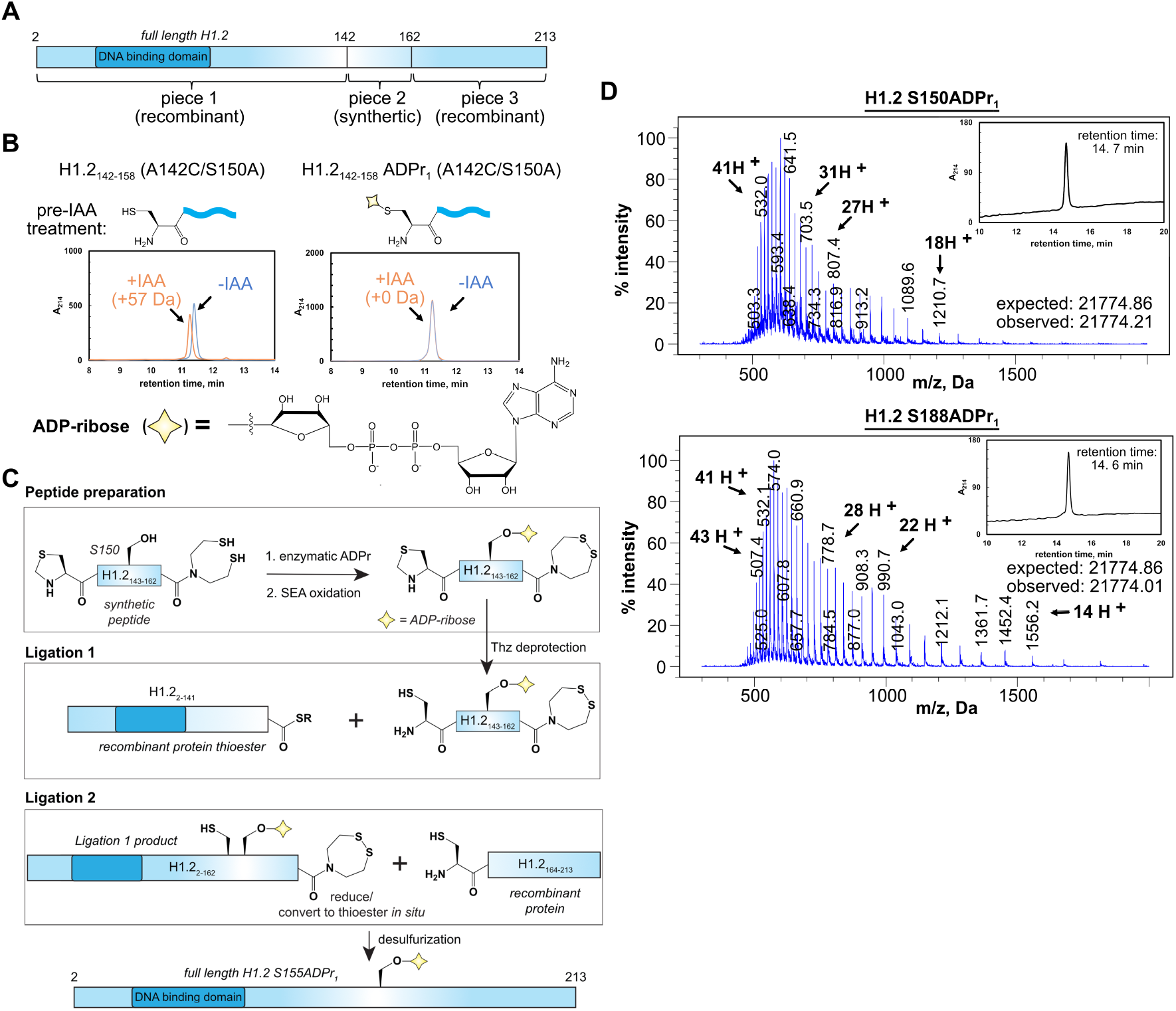
Semi-synthetic preparation of full-length, mono-ADP-ribosylated H1.2 constructs. (A) The three-piece protein assembly strategy to access the H1.2 S150ADPr site. (B) RP-HPLC analysis of the unmodified and ADP-ribosylated H1.2_142-158_ (A142C/S150A) peptides pre- and post-iodoacetamide (IAA) treatment. The mass difference post-IAA treatment is indicated. For raw ESI-MS analyses, see Figure S3. (C) Schematic summarizing the semisynthetic route to prepare full-length H1.2 S150ADPr_1_. For similar schematic depicting full-length H1.2 S150ADPr_1_ semisynthetic route, see Figure S4. (D) RP-HPLC and ESI-MS analysis of the full-length H1.2 S150ADPr_1_ and full-length H1.2 S188ADPr_1_ proteins.

Our discovery that PARP1:HPF1 efficiently modifies N-terminal cysteine side chains is a critical point to consider when preparing ADP-ribosylated proteins via the chemoenzymatic ADPr strategy. All ADP-ribosylated fragments that require an N-terminal cysteine for downstream ligation reactions must maintain a thiol protecting group through the enzymatic ADPr step. Furthermore, the peptide:ADP-ribose linkage must remain stable under conditions required to liberate the free thiol after the peptide ADPr reaction. Considering that serine ADPr is an acid-stable modification, an N-terminal thiazolidine (Thz) protection strategy was pursued, as this moiety can be rapidly and quantitatively converted to cysteine in the presence of methoxyamine hydrochloride under acidic conditions^24^. Initial N-terminal Thz H1.2_142-162,_ synthesis efforts revealed that the Thz moiety is not stable during the acyl hydrazide to thioester conversion reaction, as previously reported^25^. We therefore prepared H1.2_142-162_ as a C-terminal bis(2-sulfanylethyl)amido (SEA) peptide (Figure 2C and S2). While the oxidized SEA group is inert in native chemical ligation reactions, the reduced SEA moiety is highly reactive with small molecule thiols to generate a C-terminal thioester. Thus, the user can selectively deprotect the free N-terminal Thz or the oxidized SEA group (via incubation with 10 mM TCEP) depending on the desired sequential ligation directionality. More importantly, we found that both deprotection strategies are compatible with the ADP-ribose moiety.

ADP-ribosylated H1.2_142-162_ was prepared by incubating the Thz/reduced SEA peptide with the PARP1:HPF1 complex, and the mono-ADP-ribosylated product purified via RP-HPLC (Figure S2). Next, the peptide was resuspended in a mild oxidation buffer to fully oxidize the SEA moiety, and the Thz deprotection reaction was performed to liberate the N-terminal cysteine (Figure S2). A native chemical ligation reaction with H1.2_2-141_ was carried out and the ligated product (H1.2_2-162_ S150ADPr_1_) purified via RP-HPLC (Figure S2). An SEA to thioester conversion step was used to generate the H1.2_2-162_ S150ADPr_1_ thioester fragment, which was directly employed in a second ligation reaction with H1.2_163-213_ (Figure S2). Finally, a desulfurization reaction was carried out to convert ligation junction cysteines to native alanine residues and obtain the full-length H1.2 S150ADPr_1_ construct (Figure 2D). A similar synthetic scheme was effective to produce the full-length H1.2 S188ADPr_1_ construct (Figure 2E and S4). In this case, the synthetic fragment (amino acids 177-213) was prepared with an N-terminal Thz and C-terminal acid moiety, as a two-piece ligation strategy was sufficient to access the S188 ADPr site (Figure S5). We therefore expect the Thz-based N-terminal cysteine protection strategy will be broadly compatible with our chemoenzymatic peptide ADPr technology and, when combined with the C-terminal SEA moiety, will grant synthetic access to many unexplored ADPr sites.

We have previously shown that PARP1 efficiently elongates ADP-ribose chains from mono-ADP-ribosylated serine residues in the absence of HPF1^12^. Indeed, this catalytic property of PARP1 is maintained on the full-length, mono-ADP-ribosylated H1.2 constructs prepared herein. We found that PARP1 catalyzes ADP-ribose chain polymerization from both the S150ADPr_1_ and S188ADPr_1_ sites to a similar extent in unlabeled NAD^+^ and biotinylated NAD^+^-based ADPr assays (Figure 3A and 3B). Importantly, the ADPr activity observed in PARP1 elongation assays represents single ADP-ribose chains emanating from the pre-installed H1.2 mono-ADPr site, as only trace levels of activity could be detected on the unmodified H1.2 substrate.

**Figure 3.**
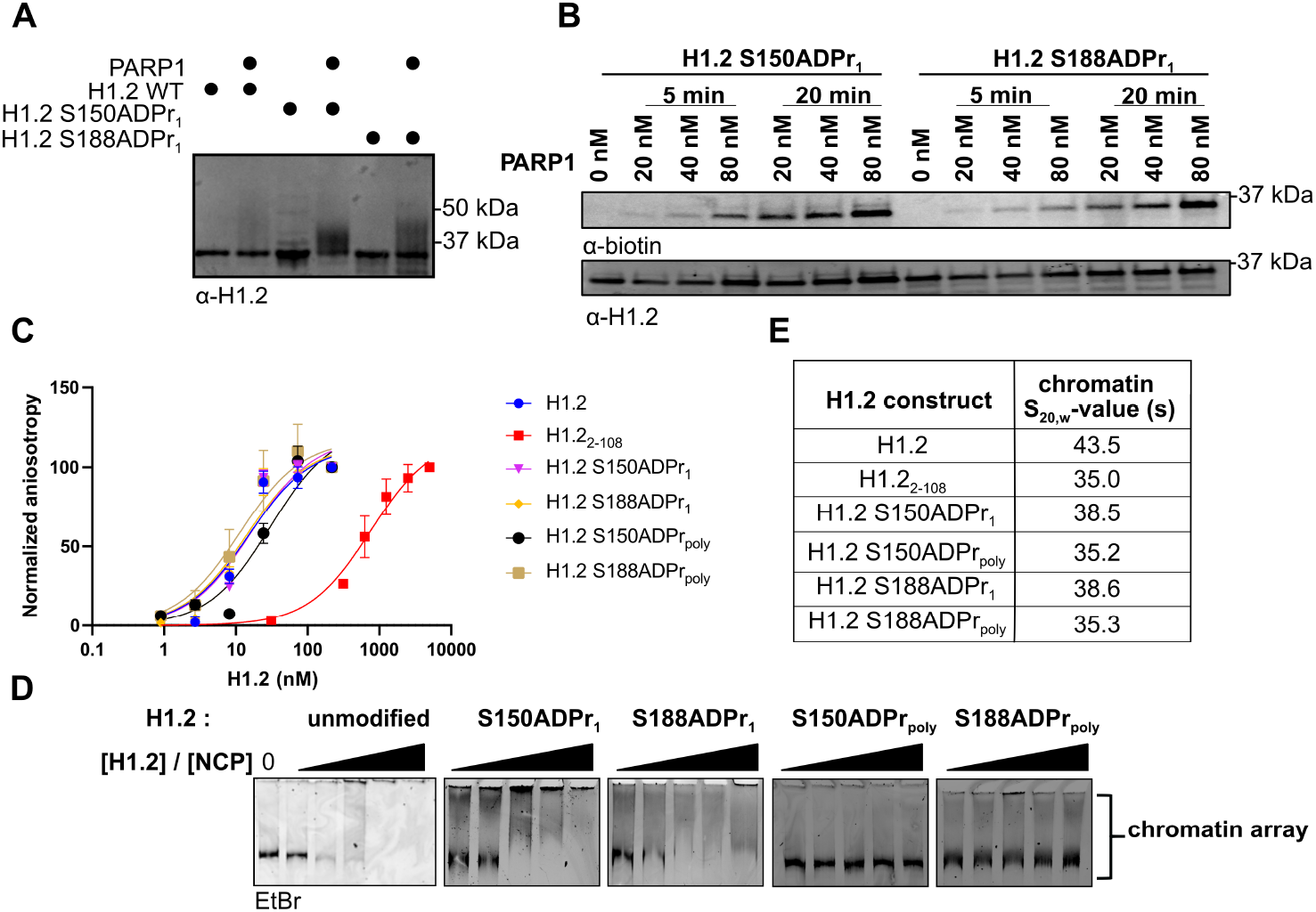
Site-specific linker histone serine ADP-ribosylation directly abrogates its ability to induce chromatin compaction. (A) Western blot analysis of PARP1 elongation activity on the indicated unmodified or mono-ADP-ribosylated full-length H1.2 substrate protein. (B) Western blot analysis of PARP1 elongation activity time course on the indicated mono-ADP-ribosylated full-length H1.2 substrate in the presence of biotinylated NAD^+^. (C) Fluorescence polarization assays to evaluate binding affinities of the indicated H1.2 constructs to a fluorescein-labeled 30 base pair DNA fragment. (D) Native 3% TBE gel electrophoresis analysis of chromatin arrays (5 nM array = 60 nM nucleosome) in the presence of increasing concentrations of the indicated H1.2 construct (120, 240, 480, 960 and 1200 nM). (E) Sedimentation coefficient values (S_20,w_) of chromatin arrays (15 nM array = 180 nM nucleosome) in the presence of the indicated H1.2 construct (180 nM) as determined via analytical ultracentrifugation.

The linker histone C-terminal domain is known to enhance DNA binding affinity^26^. A fluorescence polarization-based 30-mer DNA binding assay was employed to determine if site-specific serine mono-ADPr impacts the H1:DNA interaction (Figure 3C and Table S1). While an H1.2 construct lacking the C-terminal domain (H1.2_2-108_) exhibited a 50-fold reduction in DNA binding affinity, mono-ADPr did not significantly impact the H1.2:DNA interaction. We next sought to test the possibility that H1.2 poly-ADPr is required to disrupt the H1.2:DNA interface. To this end, the full-length H1.2 S150ADPr_1_ and H1.2 S188ADPr_1_ constructs were incubated with PARP1 and NAD^+^, and products bearing variable length ADP-ribose chains were isolated via RP-HPLC fractionation. Gel migration and mass analysis of the poly-ADP-ribosylated H1 products showed a chain length distribution ranging primarily from 4-18 ADP-ribose units (Figure S6), and the purified constructs (H1.2 S150ADPr_poly_ and H1.2 S188ADPr_poly_) were employed in DNA interaction assays. Again, no significant impact on DNA affinity was observed. While these results suggest that H1.2 poly-ADPr does not directly impact DNA binding activity, it should be noted that longer ADP-ribose chain lengths and/or chromatin templates may be required to elicit such an effect.

We next explored the effects of site-specific mono- and poly-ADPr on H1-induced chromatin compaction. Chromatin array substrates (5 nM) comprising 12 evenly spaced nucleosomes on a single DNA template were incubated with increasing concentrations of the unmodified, mono- and poly-ADP-ribosylated H1 constructs (Figure 3D). Native gel shift-based compaction assays revealed that unmodified H1 constructs abrogated gel migration of all array species at 240 nM (4:1 molar ratio of H1:nucleosome), indicating a strong compaction effect. In stark contrast, no chromatin compaction was observed in the presence of the poly-ADP-ribosylated H1 constructs at concentrations as high as 600 nM (10:1 molar ratio of H1.2:nucleosome). A more modest effect was observed with mono-ADP-ribosylated H1 constructs, which maintained their ability to stimulate chromatin compaction albeit at higher concentrations relative to the unmodified protein. Thus, site-specific H1.2 serine ADPr is sufficient to abrogate its chromatin-compaction activity, and this effect is amplified by ADP-ribose chain elongation.

To further characterize H1.2 ADPr and its impact on chromatin structure, H1.2 constructs were incubated with chromatin arrays at a 12:1 molar ratio (an equimolar ratio of H1.2:nucleosome) and chromatin sedimentation velocities were analyzed via analytical ultracentrifugation (Figure 3E and Table S2). Consistent with gel mobility results, the chromatin arrays that were incubated with unmodified H1.2 displayed the highest sedimentation coefficient (s_20,w_ = 43.5 S) and thus the greatest level of compaction. Again, we found that mono-ADPr at either the S150 or S188 site was sufficient to reduce chromatin compaction levels as evidenced by the reduced sedimentation coefficients (s_20,w_ = 38.5 S and 38.6 S, respectively) relative to the unmodified H1.2-treated arrays. These compaction deficiencies were even more pronounced in array samples treated with poly-ADP-ribosylated H1 constructs, wherein H1.2 S150ADPr_poly_ and H1.2 S188ADPr_poly_ s_20,w_ values were 35.2 S and 35.3 S, respectively. Interestingly, compaction levels measured in the presence of H1.2 S150ADPr_poly_, H1.2 S188ADPr_poly_, or the H1.2_2-108_ construct lacking the C-terminal domain were nearly identical. Therefore, poly-ADPr at a single serine site is sufficient to prevent the H1.2 C-terminal disordered domain from contributing to chromatin compaction.

In closing, we have developed a strategy to prepare ADP-ribosylated peptides that can be assembled into full-length proteins using sequential native chemical ligation reactions. We found that site-specific serine mono-ADPr on the linker histone H1.2 is sufficient to induce chromatin decompaction, and that this effect is amplified by ADP-ribose chain elongation. Importantly, our approach allows us to unambiguously attribute these chromatin compaction deficiencies to H1.2 ADPr. The results presented here are consistent with a model wherein H1.2 serine ADPr induces chromatin relaxation, but additional factors are necessary to displace H1.2 from DNA damage sites. More broadly, our work demonstrates the utility of an expanded semisynthetic toolkit to study site-specific serine ADPr and its impact on protein function and genome structure.

## Acknowledgements

We thank Dr. Deepak Nijhawan, Dr. Benjamin Tu and members of the Liszczak laboratory for insightful discussions. We thank Dr. Andrew Lemoff and the UT Southwestern Proteomics Core for technical assistance. This work was supported by grants from the Welch Foundation (I-2039-20200401 to G.L.), the Cancer Prevention Research Institute of Texas (RR180051 to G.L.) and the American Cancer Society (UTSW-IRG-17-174-13). G.L. is the Virginia Murchison Linthicum Scholar in Medical Research.

## Declaration of interests

The authors declare no competing interests.

## Supporting information

A detailed description of the materials and methods used in this study can be found in the Supporting Information. Figure S1-S6 contain MS and RP-HPLC characterizations of protein fragments and full-length proteins described in the study, as well as a schematic of the two-piece assembly strategy used to access the H1.2 S188ADPr site. Tables S1 and S2 show full data from fluorescence polarization experiments and analytical ultracentrifugation, respectively. Uncropped gels and blots presented in this study are also included here.

## Notes

### Competing Interest Statement

The authors have declared no competing interest.

## References

1 Gupte, R., Liu, Z. & Kraus, W. L. PARPs and ADP-ribosylation: recent advances linking molecular functions to biological outcomes. Genes Dev 31, 101–126, (2017).

2 Gibbs-Seymour, I., Fontana, P., Rack, J. G. M. & Ahel, I. HPF1/C4orf27 Is a PARP-1-Interacting Protein that Regulates PARP-1 ADP-Ribosylation Activity. Mol Cell 62, 432–442, (2016).

3 Bonfiglio, J. J., Fontana, P., Zhang, Q., Colby, T., Gibbs-Seymour, I., Atanassov, I., Bartlett, E., Zaja, R., Ahel, I. & Matic, I. Serine ADP-Ribosylation Depends on HPF1. Mol Cell 65, 932–940 e936, (2017).

4 Larsen, S. C., Hendriks, I. A., Lyon, D., Jensen, L. J. & Nielsen, M. L. Systems-wide Analysis of Serine ADP-Ribosylation Reveals Widespread Occurrence and Site-Specific Overlap with Phosphorylation. Cell Rep 24, 2493–2505 e2494, (2018).

5 Palazzo, L., Leidecker, O., Prokhorova, E., Dauben, H., Matic, I. & Ahel, I. Serine is the major residue for ADP-ribosylation upon DNA damage. Elife 7, (2018).

6 Rudolph, J., Roberts, G., Muthurajan, U. M. & Luger, K. HPF1 and nucleosomes mediate a dramatic switch in activity of PARP1 from polymerase to hydrolase. Elife 10, (2021).

7 Suskiewicz, M. J., Zobel, F., Ogden, T. E. H., Fontana, P., Ariza, A., Yang, J. C., Zhu, K., Bracken, L., Hawthorne, W. J., Ahel, D., Neuhaus, D. & Ahel, I. HPF1 completes the PARP active site for DNA damage-induced ADP-ribosylation. Nature 579, 598–602, (2020).

8 Hendriks, I. A., Buch-Larsen, S. C., Prokhorova, E., Elsborg, J. D., Rebak, A., Zhu, K., Ahel, D., Lukas, C., Ahel, I. & Nielsen, M. L. The regulatory landscape of the human HPF1- and ARH3-dependent ADP-ribosylome. Nat Commun 12, 5893, (2021).

9 Leidecker, O., Bonfiglio, J. J., Colby, T., Zhang, Q., Atanassov, I., Zaja, R., Palazzo, L., Stockum, A., Ahel, I. & Matic, I. Serine is a new target residue for endogenous ADP-ribosylation on histones. Nat Chem Biol 12, 998–1000, (2016).

10 Liszczak, G., Diehl, K. L., Dann, G. P. & Muir, T. W. Acetylation blocks DNA damage-induced chromatin ADP-ribosylation. Nat Chem Biol 14, 837–840, (2018).

11 Langelier, M. F., Billur, R., Sverzhinsky, A., Black, B. E. & Pascal, J. M. HPF1 dynamically controls the PARP1/2 balance between initiating and elongating ADP-ribose modifications. Nat Commun 12, 6675, (2021).

12 Mohapatra, J., Tashiro, K., Beckner, R. L., Sierra, J., Kilgore, J. A., Williams, N. S. & Liszczak, G. Serine ADP-ribosylation marks nucleosomes for ALC1-dependent chromatin remodeling. Elife 10, (2021).

13 Prokhorova, E. et al. Unrestrained poly-ADP-ribosylation provides insights into chromatin regulation and human disease. Mol Cell 81, 2640–2655 e2648, (2021).

14 Curtin, N. J. & Szabo, C. Poly(ADP-ribose) polymerase inhibition: past, present and future. Nat Rev Drug Discov 19, 711–736, (2020).

15 Bonfiglio, J. J., Leidecker, O., Dauben, H., Longarini, E. J., Colby, T., San Segundo-Acosta, P., Perez, K. A. & Matic, I. An HPF1/PARP1-Based Chemical Biology Strategy for Exploring ADP-Ribosylation. Cell 183, 1086–1102 e1023, (2020).

16 Voorneveld, J., Rack, J. G. M., Ahel, I., Overkleeft, H. S., van der Marel, G. A. & Filippov, D. V. Synthetic alpha- and beta-Ser-ADP-ribosylated Peptides Reveal alpha-Ser-ADPr as the Native Epimer. Org Lett 20, 4140–4143, (2018).

17 Hananya, N., Daley, S. K., Bagert, J. D. & Muir, T. W. Synthesis of ADP-Ribosylated Histones Reveals Site-Specific Impacts on Chromatin Structure and Function. J Am Chem Soc 143, 10847–10852, (2021).

18 Li, Z. et al. Destabilization of linker histone H1.2 is essential for ATM activation and DNA damage repair. Cell Res 28, 756–770, (2018).

19 Strickfaden, H., McDonald, D., Kruhlak, M. J., Haince, J. F., Th’ng, J. P. H., Rouleau, M., Ishibashi, T., Corry, G. N., Ausio, J., Underhill, D. A., Poirier, G. G. & Hendzel, M. J. Poly(ADP-ribosyl)ation-dependent Transient Chromatin Decondensation and Histone Displacement following Laser Microirradiation. J Biol Chem 291, 1789–1802, (2016).

20 Althaus, F. R. Poly ADP-ribosylation: a histone shuttle mechanism in DNA excision repair. J Cell Sci 102 (Pt 4), 663–670, (1992).

21 Azad, G. K., Ito, K., Sailaja, B. S., Biran, A., Nissim-Rafinia, M., Yamada, Y., Brown, D. T., Takizawa, T. & Meshorer, E. PARP1-dependent eviction of the linker histone H1 mediates immediate early gene expression during neuronal activation. J Cell Biol 217, 473–481, (2018).

22 Poirier, G. G., de Murcia, G., Jongstra-Bilen, J., Niedergang, C. & Mandel, P. Poly(ADP-ribosyl)ation of polynucleosomes causes relaxation of chromatin structure. Proc Natl Acad Sci U S A 79, 3423–3427, (1982).

23 Zheng, J. S., Tang, S., Qi, Y. K., Wang, Z. P. & Liu, L. Chemical synthesis of proteins using peptide hydrazides as thioester surrogates. Nat Protoc 8, 2483–2495, (2013).

24 Bang, D., Pentelute, B. L. & Kent, S. B. Kinetically controlled ligation for the convergent chemical synthesis of proteins. Angew Chem Int Ed Engl 45, 3985–3988, (2006).

25 Fang, G. M., Wang, J. X. & Liu, L. Convergent chemical synthesis of proteins by ligation of peptide hydrazides. Angew Chem Int Ed Engl 51, 10347–10350, (2012).

26 White, A. E., Hieb, A. R. & Luger, K. A quantitative investigation of linker histone interactions with nucleosomes and chromatin. Sci Rep 6, 19122, (2016).

