## supplemental files for "A protein semisynthesis-based strategy to probe the impact of site-specific serine ADP-ribosylation on linker histone function"

**Molecular cloning**
Standard gene cloning and mutagenesis strategies were used to generate expression constructs for all recombinant proteins. The full length H1.2 gene was purchased from the Ultimate™ ORF Lite human cDNA collection (Life Technologies), and cloned into a pET30 vector via Gibson Assembly to produce an *E. coli* expression plasmid for the H.1.2-GyrA-6xHis construct. To generate H1.2 S150ADPr construct, three-piece ligation strategy requiring two recombinantly expressed protein fragments was employed. Piece 1 (H1.2$_{2-141}$) was fused to a C-terminal GyrA intein-6xHis tag and cloned into pET30 vector to facilitate expressed protein ligation. Piece 3 (H1.2$_{164-213}$ (A164C)) was fused with a 6xHis-SUMO tag at the N-terminus and a GryA intein at C-terminus in pET30 vector. For H1.2 S188ADPr construct generation, two-piece ligation strategy requiring one recombinantly expressed protein fragment was employed. Piece 1 (H1.2$_{2-176}$) was C-terminally fused to 6xHis-tagged GyrA intein and cloned into pET30 vector to facilitate expressed protein ligation. For mammalian expression, The H1.2 gene was fused to an N-terminal 6xHis-tag and subcloned into mammalian expression plasmid (pCMV). Plasmids for PARP1, HPF1, PARG were prepared as described previously[1].

**Recombinant protein expression and purification**
Expression plasmids for full length H1.2, H1.2$_{2-141}$, H1.2$_{163-213}$, H1.2$_{2-176}$, were transformed into Rosetta2 (DE3) cells and inoculated into 1 L of Luria Broth. Protein expression was induced with 0.5 mM IPTG at a cell OD$_{600}$ of 0.6. Expression was carried out at 16 °C for 16 h. Cells were harvested by centrifugation and disrupted via sonication in a lysis buffer containing 50 mM Tris, pH 7.5, 1 M NaCl, and 1 mM PMSF. Following centrifugation at 40,000 RCF for 30 min at 4 °C, the target protein was captured on Ni-NTA pre-equilibrated in lysis buffer. Following 1 h batch binding at 4 °C, resin was washed with lysis buffer supplemented with 25 mM imidazole and protein was eluted in lysis buffer supplemented with 300 mM imidazole. For Piece 1$_{2-141}$ and Piece 1$_{2-176}$, 300 mM MES-Na and 10 mM TCEP were added at pH 7.0, and incubated at room temperature for 16 h to achieve thiolysis. For Piece 3$_{163-213}$ and full length H1.2, Ulp1 enzyme was added to the protein solution for 1 h to cleave the SUMO tag and 200 mM BME was added for 30 min at 37°C to achieve GryA fusion hydrolysis. Following thiolysis, protein solution was supplemented with 1% TFA and was spun at 4,000 × g for 30 min to remove precipitation and purified by RP-HPLC (C18 preparative column). Fractions were analyzed on analytical C18 RP-HPLC and ESI-MS and pure product (>95%) were pooled, lyophilized and stored at -80 °C.

**Recombinant H1.2 protein and protein fragments**
H1.2$_{2-213}$ (full length)
SETAPAAPAAAPPAEKAPVKKKAAKKAGGTPRKASGPPVSELITKAVAASKERSGVSLA
ALKKALAAAGYDVEKNNSRIKLGLKSLVSKGTLVQTKGTGASGSFKLNKKAASGEAKPKV
KKAGGTKPKKPVGAAKKPKKAAGGATPKKSAKKTPKKAKKPAAATVTKKVAKSPKKAKVA
KPKKAAKSAAKAVKPKAAKPKVVKPKKAAPKKK

Three-piece ligation Piece 1, H1.2 $_{2-141}$ (thioester)
SETAPAAPAAAPPAEKAPVKKKAAKKAGGTPRKASGPPVSELITKAVAASKERSGVSLAALKKA
LAAAGYDVEKNNSRIKLGLKSLVSKGTLVQTKGTGASGSFKLNKKAASGEAKPKVKKAGGTKP
KKPVGAAKKPKKA-thioester

Three-piece ligation Piece 3, H1.2 $_{163-213}$ (A163C)
CATVTKKVAKSPKKAKVAKPKKAAKSAAKAVKPKAAKPKVVKPKKAAPKKK

Two-piece ligation Piece 1, H1.2 $_{2-176}$ (thioester)

SETAPAAPAAAPPAEKAPVKKKAAKKAGGTPRKASGPPVSELITKAVAASKERSGVSLAA LKKALAAAGY DVEKNNSRIKLGLKSLVSKGTLVQTKGTGASGSFKLNKKAASGEAKPKVK KAGGTKPKKPVGAAKKPKKAAGGATPKKSAKKTPKKAKKPAAATVTKKVAKSPKK-thioester

**Immunoprecipitation of 6xHis-H1.2 constructs from mammalian cells**
293T cells were transfected using a calcium phosphate transfection kit (Takara bio) following manufacturer's protocols with plasmid DNA encoding the 6xHis-tagged linker histone (H1.2 WT, H1.2 S150A, H1.2 S188A, or H1.2 S150A/S188A). After 48 h, cells were treated with 2 mM $H_2O_2$ for 15 min, harvested in lysis buffer (8 M urea, 1000 mM NaCl, 50 mM Tris pH 7.5, 0.5% NP-40) and further disrupted using microtip with 12 % amplitude for 10 sec. Cleared lysate was incubated with pre-equilibrated Ni-NTA resin for 1 h on an end-over-end rotator at 4 °C. Resin was washed with lysis buffer and bound proteins were eluted by incubating the resin in 2X SDS loading dye and boiled at 95 °C for 10 min. Next, 20 µL of sample was loaded onto an SDS–PAGE gel (12% Tris-Glycine), and the resolved gel was transferred to a PVDF membrane for western blot analysis. Relative efficiency of transfection (α-6xHis, 66005-1-Ig, Proteintech Group) and DNA damage-induced ADP-ribosylation on His-H1.2 (α-pan-ADP-ribose, MABE1016, Millipore Sigma) were analyzed.

**Reconstituted H1.2 ADPr assay**
To determine the primary H1.2 ADPr sites, biotinylated $NAD^+$ (100 µM), PARP stimulating DNA (10 µM), full length H1.2 constructs (H1.2 WT, S150A, S188A, S150A/S188A; 50 µM), PARP1 (100 nM), and HPF1 (5 µM) were incubated in a buffer comprising 50 mM Tris pH 7.5, 20 mM NaCl, 2 mM TCEP, 2 mM $MgCl_2$ at 30 °C. Reactions were quenched after 20 min using 4X-SDS loading dye and loaded onto an SDS–PAGE gel (12% Bis–Tris), and the resolved gel was transferred to a PVDF membrane for western blot analysis. Sample loading (α-H1.2, PIPA542819, ThermoFisher Scientific) and DNA damage-induced ADPr on recombinant H1.2 (α-biotin, NC9386176, Thermo-Fisher Scientific) were analyzed.

**Peptide synthesis**
*Synthetic peptides described in this study*
H1.2 $_{143\text{-}158}$: GGATPKKSAKKTPKKA-$CONH_2$
H1.2 $_{143\text{-}158}$ (S150A): GGATPKKAAKKTPKKA-$CONH_2$
H1.2 $_{142\text{-}158}$ (A142C/S150A): CGGATPKKAAKKTPKKA-$CONH_2$
H1.2 $_{166\text{-}181}$: VTKKVAKSPKKAKVAK-$CONH_2$
H1.2 $_{166\text{-}181}$ (P174A): VTKKVAKSAKKAKVAK-$CONH_2$
H1.2 $_{182\text{-}195}$: PKKAAKSAAKAVKP-$CONH_2$
H1.2 $_{182\text{-}195}$ (S188A): PKKAAKAAAKAVKP-$CONH_2$
H1.2 $_{142\text{-}162}$ (A142Thz, SEA): Thz- GGATPKKSAKKTPKKAKKPA-SEA
H1.2 $_{177\text{-}213}$ (A177Thz): KVAKPKKAAKSAAKAVKPKAAKPKVVKPKKAAPKKK- COOH

All fluorenylmethyloxycarbonyl (Fmoc)-protected amino acids were purchased from Oakwood Chemical or Combi-Blocks. Peptide synthesis resins (Trityl-OH ChemMatrix and Rink Amide ChemMatrix) were purchased from Biotage. All analytical reversed-phase HPLC (RP-HPLC) was performed on an Agilent 1260 series instrument equipped with a quaternary pump and an XBridge Peptide C18 (5 µm, 4 × 150 mm; Waters) at a flow rate of 1 mL/min. Similarly, semi-preparative scale purifications were performed employing an XBridge Peptide C18 semi-preparative column (5 µm, 10 mm × 250 mm, Waters) at a flow rate of 4 mL/min. Preparative RP-HPLC was performed on an Agilent 1260 series instrument equipped with a preparatory pump and an XBridge Peptide C18 preparatory column (10 µM; 19 × 250 mm, Waters) at a flow rate of 20 mL/min. All instruments were equipped with a variable wavelength UV-detector. All RP–HPLC steps were performed using 0.1% (trifluoroacetic acid (TFA) Oakwood Chemical) in $H_2O$

(Solvent A) and 90% acetonitrile (Sigma-Aldrich), 0.1% TFA in $H_2O$ (Solvent B) as mobile phases. For LC/MS analysis, 0.1% formic acid (Sigma-Aldrich) was substituted for TFA in mobile phases. Mass analysis was carried out for each product on an LC/MSD (Agilent Technologies) equipped with a 300SB-C18 column (3.5 μM; 4.6 × 100 mm, Agilent Technologies) or a X500B QTOF (Sciex).

The above amidated peptides were synthesized via solid-phase peptide synthesis on a CEM Discover Microwave Peptide Synthesizer (Matthews, NC) using the Fmoc-protection strategy on Rink Amide-ChemMatrix resin (0.5 mmol/g). For coupling reactions, amino acids (5 eq) were activated with N,N-diisopropylcarbodiimide (DIC, 5 eq, Oakwood Chemical)/Oxyma (5 eq, Oakwood Chemical) and heated to 90 °C for 2 min while bubbling with $N_2$ in N,N dimethylformamide (DMF, Oakwood Chemical). Fmoc deprotection was carried out with 20% piperidine (Sigma-Aldrich) in DMF supplemented with 0.1 M 1-hydroxybenzotriazole hydrate (HOBt, Oakwood Chemical) at 90 °C for 1 min while bubbling with $N_2$. Cleavage from resin was performed with 92.5% TFA, 2.5% triisopropylsilane (TIS, SigmaAldrich), 2.5% 1,2-ethanedithiol (EDT, Sigma-Aldrich), and 2.5% H2O for 2 h at 25 °C on an end-over-end rotisserie. The crude peptide was then precipitated by the addition of a 10-fold volume of ice-cold ether and centrifuged at 4,000 RCF for 10 min at 4 °C. The pellet was resuspended in solvent A and purified via preparative C18 RP-HPLC. Fractions were analyzed on analytical C18 RP-HPLC and ESI-MS and those containing pure product (>95%) were pooled, lyophilized and stored at -80 °C.

All peptides containing a C-terminal SEA moiety were synthesized similarly to the amidated peptides described above with the following modifications. Bis(2-sulfanylethyl)amido (SEA) polystyrene resin was using a protocol from Ollivier et al, 2014[2]. For manual loading of the first amino acid to SEA resin (0.2 mmol/g), Fmoc-alanine (10 eq), HATU (10 eq, Oakwood Chemical), and DIPEA (30 eq) were mixed in DMF and the resin was bubbled with $N_2$ for 1 h. This step was repeated with fresh reagents to ensure complete loading. The resin was then washed with DMF and bubbled in acetic anhydride:DIPEA (20 eq:40 eq) in DMF for 20 min to quench unreacted sites. Subsequent peptide synthesis reactions were performed on an automated, microwave synthesizer as described for amidated peptides. Resin cleavage was performed with 95% TFA, 2.5% TIS, and 2.5% $H_2O$ for 2 h at 25 °C on an end-over-end rotisserie. Purification of SEA peptides were performed as described for amidated peptides.

Peptides containing a C-terminal acid moiety were made on PEG (trityl-OH) resin (Biotage). Trityl-OH resin was bubbled with $N_2$ for 90 min in 5% thionyl chloride in DCM. The chlorination step was carried out again after washing the resin with DCM. Prior to manual loading, the resin was washed with 5% DIPEA in DMF, and Fmoc-lysine (4 eq) and DIPEA (4.5 eq) were mixed in DMF and bubbled with $N_2$ for 1 h. This step was repeated with fresh reagents to ensure complete loading. The resin was then washed with DMF and bubbled in DMF:MeOH:DIPEA (17:2:1) mixture for 20 min to quench unreacted sites. Subsequent peptide synthesis reactions were performed on an automated, microwave synthesizer as described for amidated peptides. Resin cleavage was performed with 95% TFA, 2.5% TIS, and 2.5% $H_2O$ for 2 h at 25 °C on an end-over-end rotisserie. Purification of C-terminal acid peptides were performed as described for amidated peptides.

**Preparation of homogenous mono-ADP-ribosylated peptides**

Peptides bearing N-terminal thiazolidine and C-terminal SEA or acid (0.5 mM) were incubated with recombinant PARP1 (1 μM), HPF1 (25 μM), and PARG (5 μM) proteins in a buffer containing 50 mM Tris pH 7.5, 20 mM NaCl, 2 mM TCEP, 2 mM $MgCl_2$, 10 mM $NAD^+$,10 μM

PARP stimulating DNA 20 min at 30 °C. Reactions were quenched with 6 M guanidine-HCl, 0.1 M sodium phosphate and were purified on a preparative C18 RP-HPLC column and pure fractions were pooled, lyophilized, and stored at -80 °C. Purified, mono-ADP-ribosylated peptides were carried forward as described in 'Assembly of H1.2 S150ADPr$_1$' and 'Assembly of H1.2 S188ADPr$_1$' methods sections.

**Iodoacetamide labeling of peptides**

H1.2$_{142-158}$ (A142C/S150A) or H1.2$_{142-158}$ ADPr$_1$ (C142/S150A) peptides (0.5 mM) were incubated with 20 mM iodoacetamide in a buffer containing 50 mM Tris pH 7.5, 20 mM NaCl, 2 mM $MgCl_2$ at 25 °C. After 1 h in the dark, the reaction mixture was quenched with solvent A and was immediately analyzed via C18 RP-HPLC and ESI-MS.

**Assembly of H1.2 S150ADPr$_1$**

For mono-ADP-ribosylated peptide preparation, see 'Preparation of homogenous mono-ADP-ribosylated peptides'. Prior to ligation, the C-terminal SEA functionality was oxidized by incubating 10 mM peptide in buffer containing 6 M guanidine-HCl, 0.1 M sodium phosphate and 5% (v/v) DMSO at pH 7.5-8.0. This reaction was nutated at 25 °C for 16 h and SEA ring oxidation was confirmed by a -2 Da mass change via LC-MS. Once quantitative oxidation was complete, 200 mM of O-methylhydroxylamine HCl (Combi-Blocks) was added to the reaction mixture. The reaction pH was then adjusted to 4.0 and incubated at 37 °C for 1 h. Deprotection of the thiazolidine ring was confirmed by a -12 Da mass change via LC-MS. H1.2$_{2-141}$ (bearing MES thioester; ~5 mM) and H1.2$_{142-162}$ S150ADPr$_1$ (bearing free N-terminal cysteine and oxidized SEA ring; ~8 mM) were combined in a degassed buffer of 6 M guanidine-HCl, 0.1 M sodium phosphate, and 100 mM 2,2,2-trifluoroethanethiol (TFET, SigmaAldrich) and adjusted to pH 7.0. Excess H1.2$_{142-162}$ was used to ensure quantitative conversion of peptide is to ligated product, which was important for efficient separation of product and starting material during purification. Reactions were incubated at 37 °C for 4 h and progress was monitored via C18 RP-HPLC and ESI-MS analysis. Note: reducing agents are omitted from this ligation step and the C-terminal oxidized SEA ring remained intact and inert throughout the reaction. Ligation products were purified on C18 RP-HPLC column and fractions containing target mass were pooled, lyophilized, and stored at -80 °C. This product is referred to as H1.2$_{2-162}$ S150ADPr$_1$. The lyophilized H1.2$_{2-162}$ S150ADPr$_1$ construct was resuspended in 0.3-1 ml of degassed buffer containing 6 M guanidine-HCl, 0.1 M sodium phosphate, 200 mM MES-Na (Sigma-Aldrich), and 50 mM TCEP and adjusted to pH 4.0 (H1.2$_{2-162}$ S150ADPr$_1$ construct concentration = ~1.5 mM). This reaction was incubated at 37 °C for 16 h to convert the C-terminal SEA moiety to a MES-thioester as confirmed by a -5 Da mass shift via LC-MS. Once MES conversion was complete, H1.2$_{163-213}$ (bearing an N-terminal cysteine; ~0.5 mM) was directly added to this reaction mixture. Reactions were supplemented with 20 mM TCEP and 100 mM TFET, adjusted to pH 7.0, and incubated 37 °C for 16 h. Excess H1.2$_{2-162}$ S150ADPr$_1$ was used to ensure quantitative consumption of H1.2$_{163-213}$, which was important for efficient separation of product from starting material during purification. Upon completion of the ligation reaction as judged by RP-HPLC and ESI-MS analysis, the reaction mixture was dialyzed against a buffer containing 6 M guanidine-HCl and 0.1 M sodium phosphate at pH 4.0 for 3 h. Desulfurization of the final product to the native H1.2 sequence was achieved through free radical-mediated desulfurization by addition of 200 mM TCEP, 60 mM VA-044 radical initiator and 150 mM of reduced glutathione. Reactions were adjusted to pH 7.0 and incubated at 37 °C for 16 h. Ligation products were purified on a semi-preparative C18 RP-HPLC column and pure fractions were pooled, lyophilized, and stored at -80 °C. Final purities of >95% were judged by analytical RP-HPLC, ESI-MS analysis and immunoblot analysis.

**Assembly of H1.2 S188ADPr$_1$**

For mono-ADP-ribosylated peptide preparation, see ‘Preparation of homogenous mono-ADP-ribosylated peptides’. H1.2 $_{177\text{-}213}$ S188ADPr$_1$ (A177Thz; 10 mM) was resuspended in buffer containing 6 M guanidine-HCl, 0.1 M sodium phosphate, and 200 mM O-methylhydroxylamine HCl, and the reaction pH was then adjusted to 4.0. The reaction was incubated at 37 °C for 1 h and deprotection of the thiazolidine ring was confirmed by a -12 Da mass change via LC-MS. The reactions were purified on a preparative C18 RP-HPLC column and pure fractions were pooled, lyophilized, and stored at -80 °C. H1.2$_{2\text{-}176}$ (bearing MES thioester; ~10 mM) and H1.2 $_{177\text{-}213}$ S188ADPr$_1$ (A177C; bearing free N-terminal cysteine and C-terminal acid moiety; ~8 mM) were combined in a degassed buffer of 6 M guanidine-HCl, 0.1 M sodium phosphate, and 100 mM 2,2,2-trifluoroethanethiol (TFET, SigmaAldrich) and adjusted to pH 7.0. Reactions were incubated at 37 °C for 4 h and progress was monitored via C18 RP-HPLC and ESI-MS analysis. Upon completion of the ligation reaction as judged by RP-HPLC and ESI-MS analysis, the reaction mixture was dialyzed against a buffer containing 6 M guanidine-HCl and 0.1 M sodium phosphate at pH 4.0 for 3 h. Desulfurization of the final product to the native H1.2 sequence was achieved through free radical-mediated desulfurization by addition of 200 mM TCEP, 60 mM VA-044 radical initiator and 150 mM of reduced glutathione. The pH of reaction mixture was adjusted to 7.0 and incubated at 37 °C for 16 h. Ligation product was purified on semi-preparative C18 RP-HPLC column and pure fractions were pooled, lyophilized, and stored at -80 °C. Final purities of >95% were judged by analytical RP-HPLC, ESI-MS analysis and immunoblot analysis.

**Poly-ADPr assays and preparation of H.1.2 S150ADPr$_{poly}$ and H1.2 S188ADPr$_{poly}$**

Poly-ADPr H1.2 constructs were prepared as described in ‘Peptide mono/poly ADPr preparation’ with following modifications. 50 µM of H1.2 WT, H1.2 S150ADPr$_1$, or H1.2 S188ADPr$_1$ was incubated with 1 µM PARP1, 10 µM of PARP stimulating DNA, 10 mM $NAD^+$ in 50 mM Tris pH 7.5, 20 mM NaCl, 2 mM TCEP and 2 mM $MgCl_2$. Reactions were incubated at 30 °C for 20 min and quenched with 4X-SDS loading dye and loaded onto an SDS–PAGE gel (12% Tris-Glycine), and the resolved gel was transferred to a PVDF membrane for western blot analysis. Sample loading as well as variable length ADP-ribose chains on recombinant H1.2 (α-H1.2, PIPA542819, Thermo-Fisher Scientific) were analyzed. For HPLC purification, the reactions were quenched with 6 M guanidine-HCl, 0.1 M sodium phosphate and elongated H1.2 were isolated on semi-preparative C18 RP-HPLC column and pooled and lyophilized. Purified proteins were refolded into a buffer comprising 50 mM Tris pH 7.5, 20 mM NaCl, and 2 mM $MgCl_2$ and the protein concentrations were measured via BCA Protein Assay Kit (ThermoFisher Scientific). For PARP1 ADP-ribose chain elongation assays comprising biotin-$NAD^+$, a similar buffer was employed wherein unlabeled $NAD^+$ was replaced with biotin-$NAD^+$ (100 µM). In these assays, 0-80 µM of full length H1.2 and 1 µM PARP1 were incubated at 30 °C for 5 or 20 min and reactions were quenched via addition of 4X SDS loading dye. Samples were loaded onto an SDS–PAGE gel (12% Tris-Glycine), and the resolved gel was transferred to a PVDF membrane for western blot analysis. Sample loading (α-H1.2, PIPA542819, Thermo-Fisher Scientific) and ADP-ribose chain elongation on recombinant H1.2 (α-biotin, NC9386176, ThermoFisher Scientific) were analyzed.

**Fluorescence polarization-based DNA interaction assays**

FP assays were performed as previously described with the following modifications[1]. Fluorescently-labeled 30-mer DNA was prepared by annealing the following nucleotides:

ATCATTAATA/iFluorT/GAATTCGCCACATGCA (IDT) and
TGCATGTGGCGAATTCATATTAATGAT

at 95 °C for 5 min. The solution was then left to cool to 25 °C over 45 min in 1x Cutsmart Buffer (NEB). Annealed DNA was used at 5 nM in a buffer containing 25 mM Tris, pH 7.5, 100 mM NaCl, 2 mM $MgCl_2$, 0.001% Triton X100, and 1 mM DTT. To analyze DNA-H1.2 interaction, H1.2 was added to final concentrations ranging from 0.9 nM to 219 nM or 30 nM to 1000 nM. Reactions were added to a black, flat-bottom 96-well plate (Corning Costar) and analyzed on a BioTek Cytation five imager equipped with a Green FP filter set (excitation: 485 nm, emission: 528 nm). Polarization values were converted to anisotropy using the following formula[3]:

$$r=(2 P/(3 - P))$$

Following background subtraction and normalization, data was processed in GraphPad Prism using a non-linear regression analysis to obtain $K_{d,app}$ values for each DNA:H1.2 interaction. Error bars represent standard deviation value from three biological replicates.

**Preparation of chromatin arrays**

Chromatin arrays were prepared as previously described[4]. Briefly, core histones (H2A, H2B, H3 and H4) were purified and octamers were assembled as described previously[1]. The DNA used for assembly of the chromatin arrays was a dodecameric repeat of the 147 bp Widom's 601 sequence, separated by 30 bp linkers (12x-177-601 DNA). Chromatin array DNA was generated upon EcoRV digestion of a pWM530 plasmid (which contained the described insert)[5] and subsequently purification via size-selective precipitation using 6% PEG-8000. The purified 12x-177-601 DNA was resuspended in water to a concentration of 1 µg/µL, and ~100 pmol was added to ~150 pmol of octamer and ~35 pmol of MMTV DNA in a buffer volume of 75 µL containing 2 M KCl, 10 mM Tris, pH 7.5, 0.1 mM EDTA, 1 mM DTT at 4 °C. The mixture was added to a Slide-A-Lyzer MINI dialysis button (3.5 kDa MWCO, ThermoFisher) and dialyzed against a buffer containing 10 mM Tris (pH 7.5), 1.4 M KCl, 0.1 mM EDTA and 1 mM DTT at 4 °C for 1.5 h. To this dialysis tank, ~350 mL of End Buffer containing 10 mM Tris (pH 7.5) 10 mM KCl, 0.1 mM EDTA, 1 mM DTT was added at a rate of 1 mL/min. After 16 h, the sample was further dialyzed twice against fresh End Buffer for 2 h each. After dialysis, any precipitation was removed through centrifugation at 20,000 RCF for 10 min at 4 °C. Chromatin arrays were purified by selective precipitation using 5 mM $MgCl_2$. The precipitated arrays were then resuspended in the End Buffer to an $A_{260}$ of ~ 2.5 and stored at 4 °C.

**Chromatin array electrophoretic mobility shift assays**

For chromatin array EMSA, the histone H1.2 constructs were diluted into End buffer at different concentrations as indicated in Figure 3D. Chromatin arrays at a final concentration of 5 nM were added to the above mixtures such that the total volume was 5 µL, and incubated at room temperature for 10 min. These reactions were then run on native 3% TBE gels after supplementation with 12% sucrose and stained with ethidium bromide for visualization of the arrays.

**Analytical ultracentrifugation**

Samples for AUC analysis was prepared by dialyzing H1.2 constructs with chromatin arrays at a 12:1 molar ratio against Working Buffer (10 mM Tris pH 7.8, 10 mM KCl, 0.1 mM EDTA) overnight at 4 °C. Working Buffer was introduced into the reference sectors of a dual-sector charcoal-filled Epon centerpieces that had been situated between two sapphire windows in standard AUC cell housings. The samples were pipetted into the sample sides of the centerpieces. Approximately 410 µL of the respective solutions were pipetted into the sectors. The cells were placed into an An50-Ti 8-hole rotor, which was positioned in a Beckman-Coulter Optima XL-I centrifuge. The chamber was evacuated, and temperature equilibration at 20 °C was undertaken for approximately 2.5 h. After that, centrifugation was commenced at 20,000 rpm. Concentration-profile data were collected using the absorbance optics tuned to 260 nm. Centrifugation continued overnight. The data were analyzed using the *c*(*s*) method in SEDFIT[6]. An *s*-resolution of 150 was employed, with $s_{min}$ and $s_{max}$ set to 0.2 S and 200 S, respectively. The position of the meniscus, the frictional ratio, and time-invariant (TI) noise were fitted. A regularization level of 0.683 was employed, as was Tikhonov-Phillips ($2^{nd}$ - derivative) regularization. The partial-specific volume was assumed to 0.622 mL/g. The buffer density and buffer viscosity were calculated by SEDNTERP[7], and were, respectively, 0.99909 g/mL and 0.01004 Poise. All sedimentation coefficient distributions were corrected to the hydrodynamicists' standard of water at 20 °C.

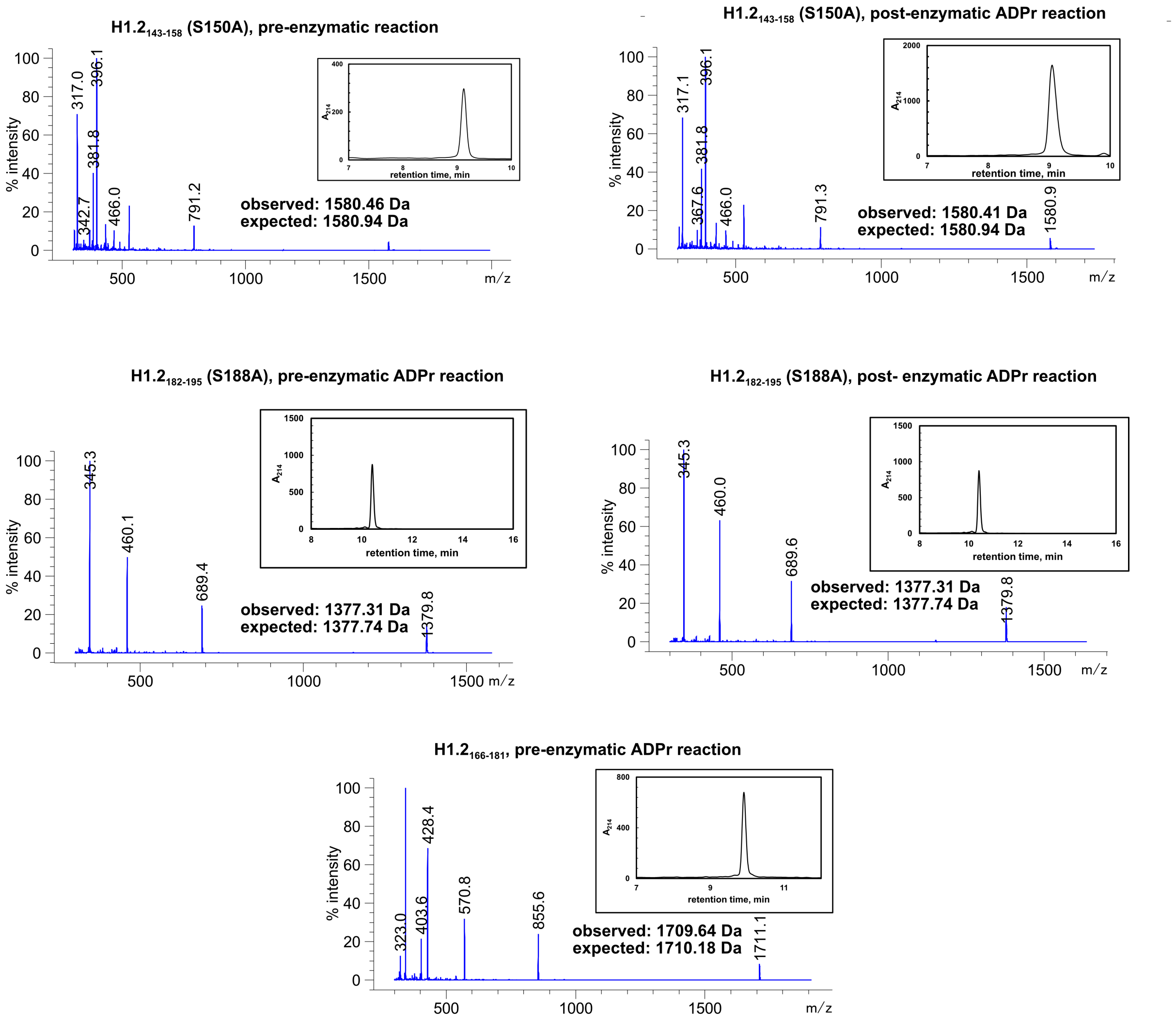


**Figure S1. ESI-MS and RP-HPLC characterizations of H1.2 peptides.** All synthetic peptides were analyzed on C18-RP-HPLC with a linear gradient from 0-30 % B over 30 min except for H1.2$_{143-158}$(S150A) which was analyzed with a linear gradient from 0-19.5% B over 15 min.

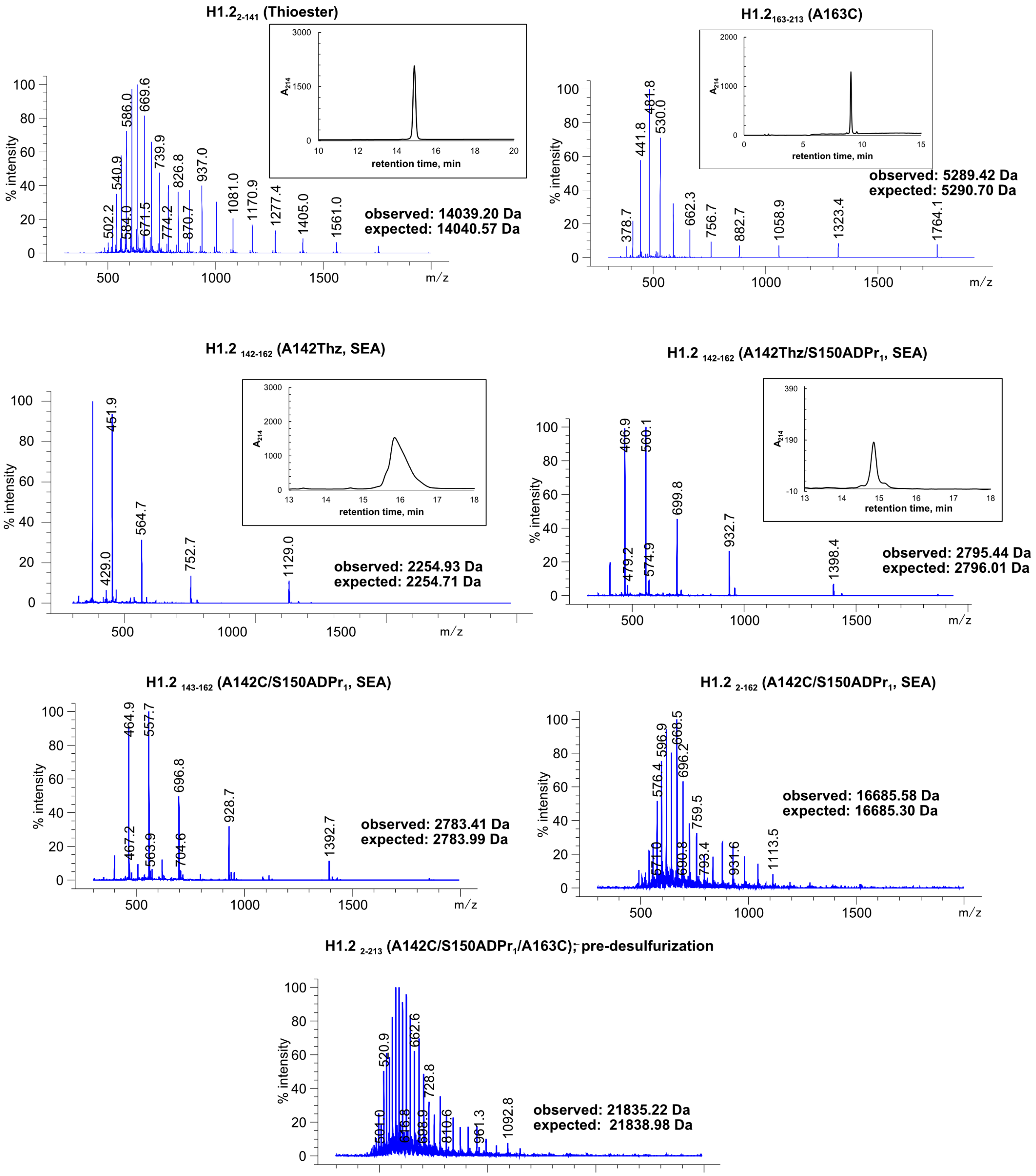


**Figure S2. ESI-MS and RP-HPLC characterizations of pieces used to assemble H1.2 S150ADPr$_1$** All synthetic peptides were analyzed on C18-RP-HPLC with a linear gradient 0-30% B over 30 min. All recombinant constructs were analyzed with a linear gradient 0-80% B over 20 min.

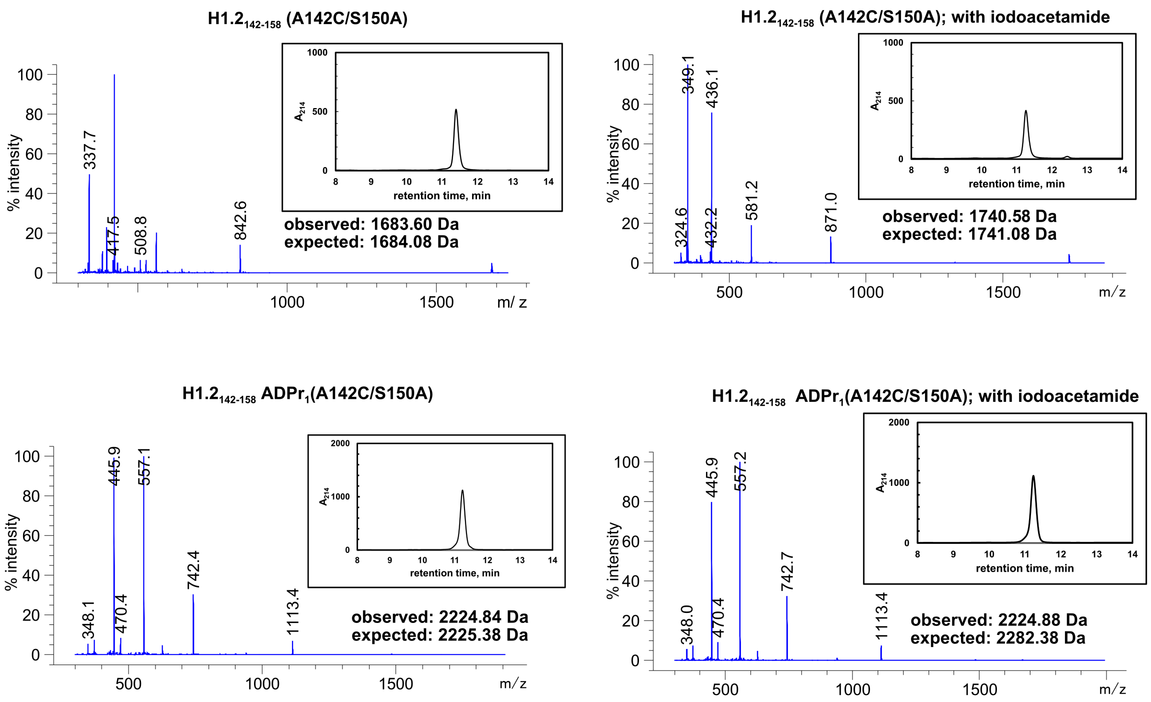


**Figure S3. ESI-MS and RP-HPLC characterizations of iodoacetamide-treated peptides.** All synthetic peptides were analyzed on C18-RP-HPLC with a linear gradient from 0-19.5% B over 15 min.

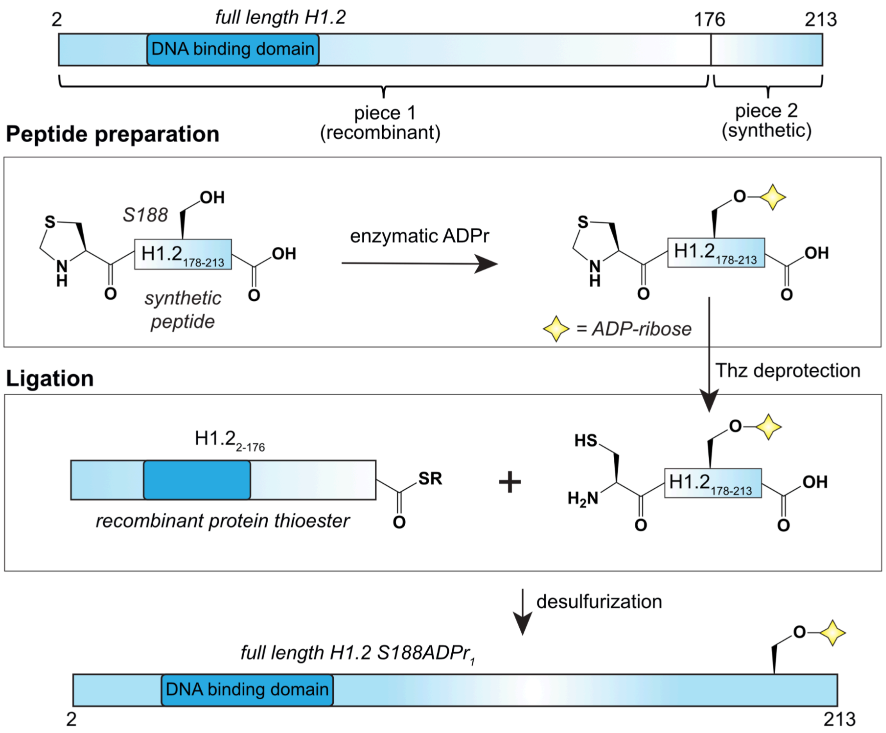


**Figure S4.The two-piece protein assembly strategy to access the H1.2 S188ADPr site.**

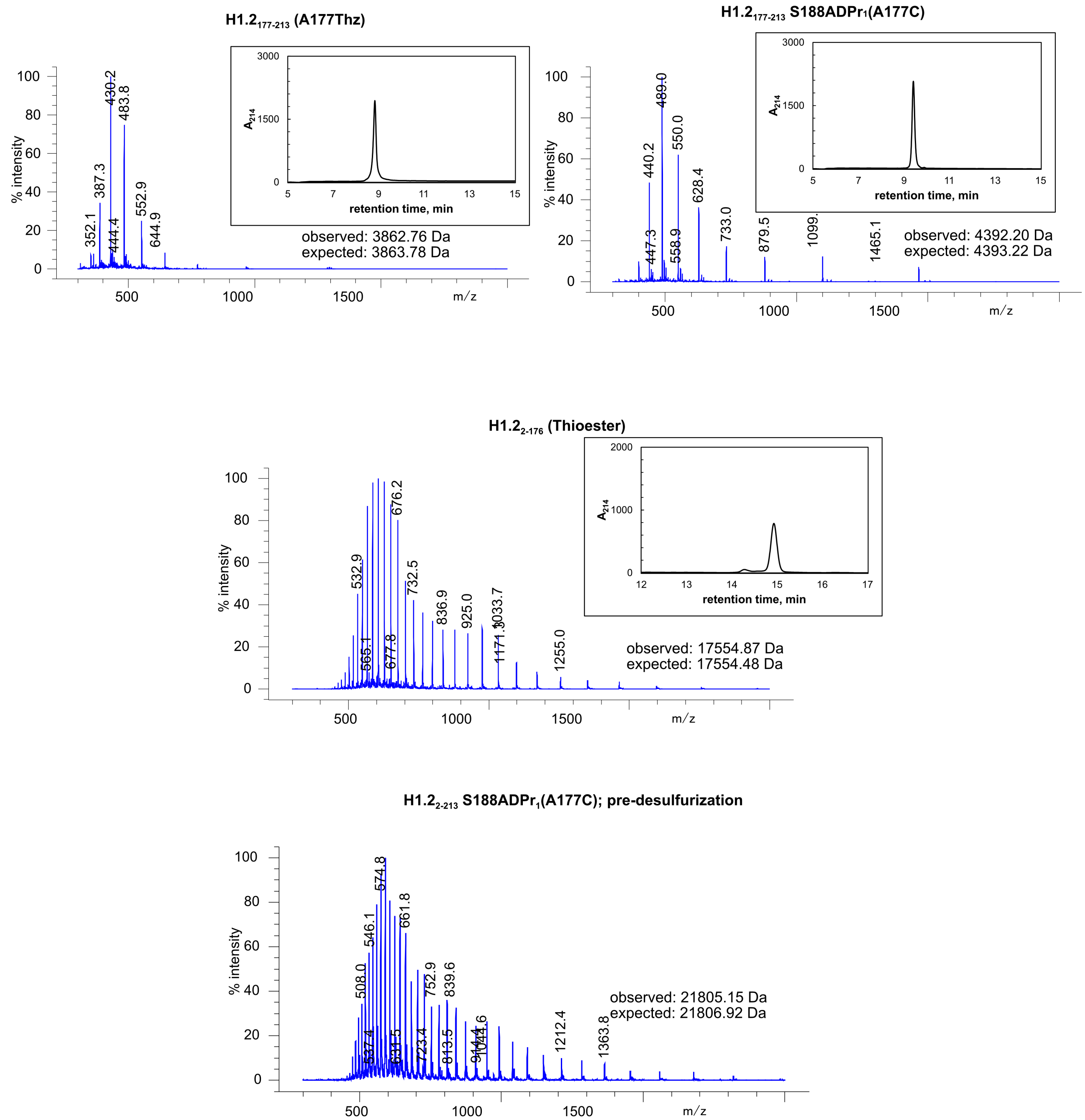


**Figure S5. ESI-MS and RP-HPLC characterizations of pieces used to assemble H1.2 S188ADPr$_1$**
All constructs were analyzed on C18-RP-HPLC with a linear gradient of 0-80% B over 20 min.

**H1.2 S150ADPr$_{poly}$**

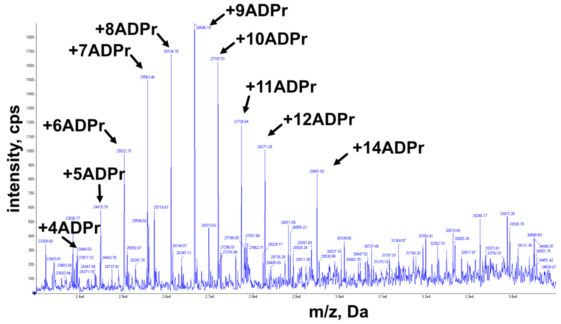


**H1.2 S188ADPr$_{poly}$**

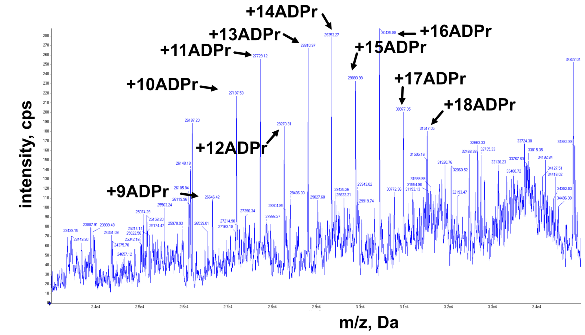


**Figure S6. Deconvolution of QTOF analysis of isolated H1.2 S150ADP$_{poly}$ species and S188ADPr$_{poly}$ species**

### Uncropped gels

#### Fig 1B

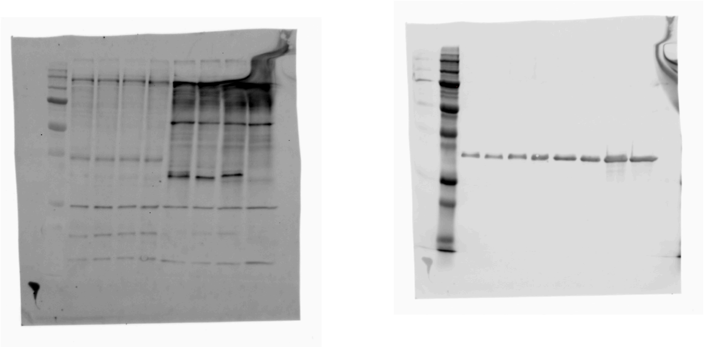

#### Fig 1C

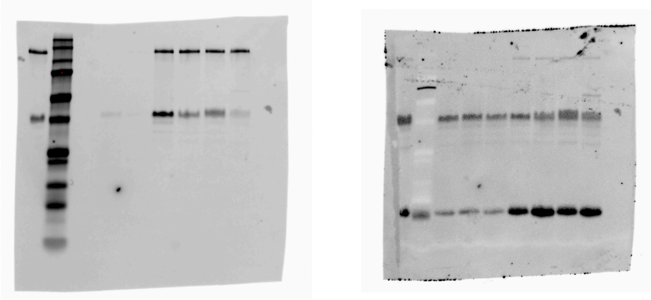

#### Fig 3A

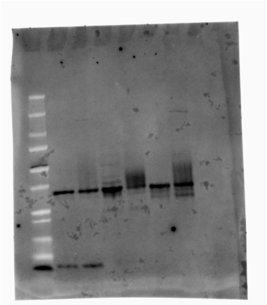

#### Fig 3B

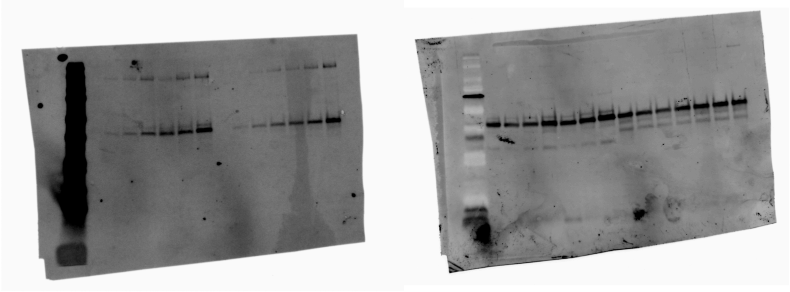

uncropped gels from Fig 1 and Fig 3A-B

**Uncropped gels**

**Fig 3D**

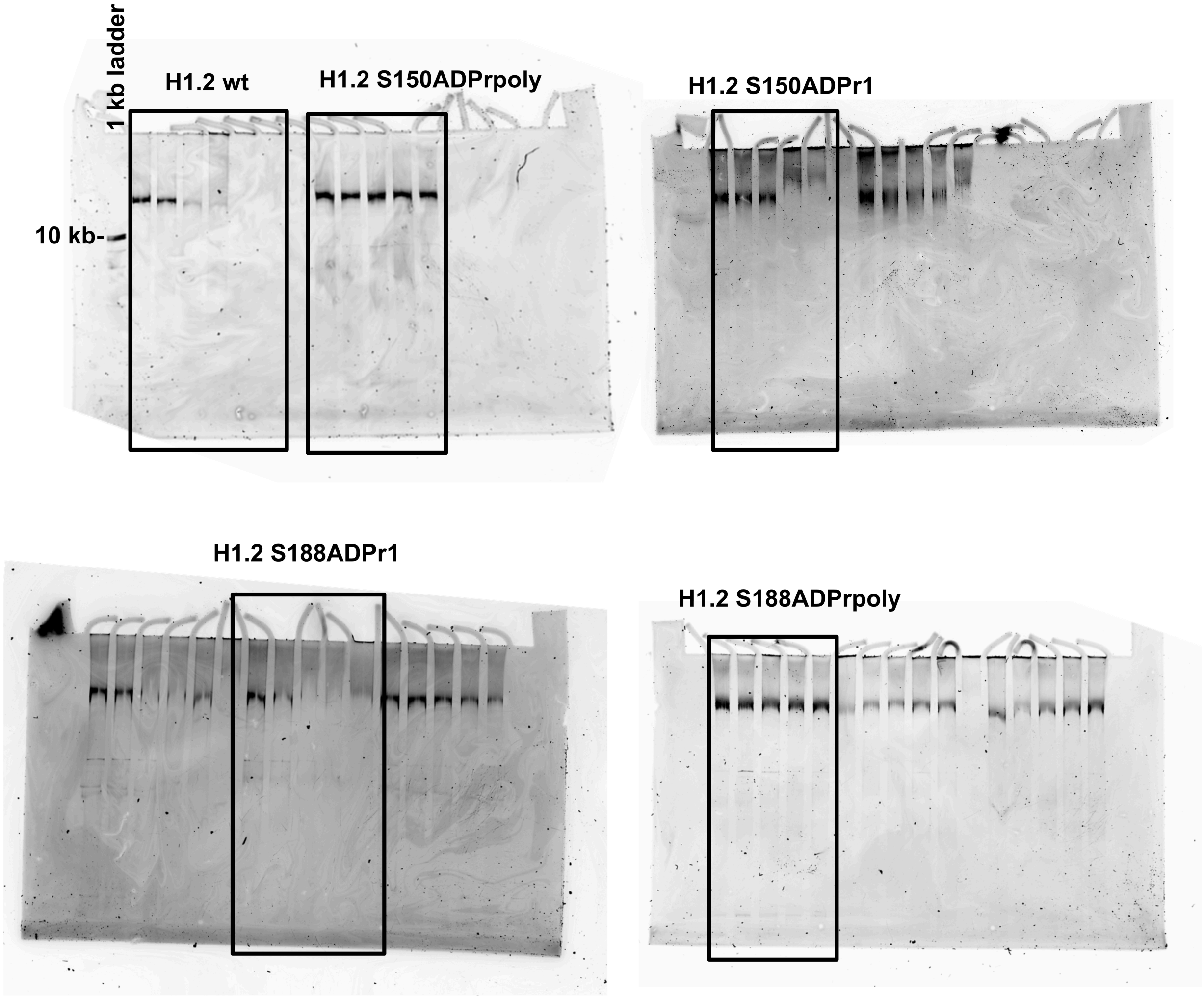


uncropped gels from Fig 3D with relevant lanes boxed

**Table S1**

| **H1.2 construct** | H1.2 | $H1.2_{2-108}$ | H1.2 $S150ADPr_1$ | H1.2 $S188ADPr_1$ | H1.2 $S150ADPr_{poly}$ | H1.2 $S188ADPr_{poly}$ |
|---|---|---|---|---|---|---|
| **$K_{d,\,app}$ (nM)** | 15.07 | 756 | 16.64 | 14.18 | 30.3 | 11.37 |
| **95% CI** | 9.897 to 22.90 | 543.7 to 1043 | 10.47 to 26.40 | 9.658 to 20.76 | 19.58 to 46.91 | 7.118 to 17.93 |
| **$R^2$** | 0.9343 | 0.9638 | 0.9195 | 0.9440 | 0.9348 | 0.9198 |

**Table S1. Fluorescence polarization data from H1.2:DNA interaction assays.** $K_{d,\,app}$ = Kd apparent CI = confidence interval, $R^2$= R-squared value

**Table S2**

| **H1 construct** | **Chromatin $S_{20,w}$ -value (s)** | **$f/f_0$** |
|---|---|---|
| H1.2 | 43.5 | 2.81 |
| $H1.2_{2-108}$ | 35.0 | 3.18 |
| H1.2 $150SADPr_1$ | 38.5 | 3.21 |
| H1.2 $150SADPr_{poly}$ | 35.2 | 3.06 |
| H1.2 $188SADPr_1$ | 38.6 | 2.86 |
| H1.2 $188SADPr_{poly}$ | 35.3 | 2.91 |

**Table S2. Sedimentation properties of the nucleosome arrays.** $f/f_0$ = frictional ratio

**References:**


1 Mohapatra, J., Tashiro, K., Beckner, R. L., Sierra, J., Kilgore, J. A., Williams, N. S. & Liszczak, G. Serine ADP-ribosylation marks nucleosomes for ALC1-dependent chromatin remodeling. *Elife* **10**, (2021).

2 Ollivier, N., Raibaut, L., Blanpain, A., Desmet, R., Dheur, J., Mhidia, R., Boll, E., Drobecq, H., Pira, S. L. & Melnyk, O. Tidbits for the synthesis of bis(2-sulfanylethyl)amido (SEA) polystyrene resin, SEA peptides and peptide thioesters. *J Pept Sci* **20**, 92-97, (2014).

3 Lakowicz, J. R. *Principles of Fluorescence Spectroscopy*. (Springer Science+Business Media, LLC, 2006).

4 Muller, M. M., Fierz, B., Bittova, L., Liszczak, G. & Muir, T. W. A two-state activation mechanism controls the histone methyltransferase Suv39h1. *Nat Chem Biol* **12**, 188-193, (2016).

5 Dorigo, B., Schalch, T., Bystricky, K. & Richmond, T. J. Chromatin fiber folding: requirement for the histone H4 N-terminal tail. *J Mol Biol* **327**, 85-96, (2003).

6 Schuck, P. Size-distribution analysis of macromolecules by sedimentation velocity ultracentrifugation and lamm equation modeling. *Biophys J* **78**, 1606-1619, (2000).

7 Laue, T. M., Shah, B. D., Ridgeway, T. M. & Pelletier, S. L. Computer-aided interpretation of analytical sedimentation data for proteins. *Analytical Ultracentrifugation in Biochemistry and Polymer Science* (eds. Harding S.E., Rowe A.J., & Horton J.C.) The Royal Society of Chemistry, Cambridge, United Kingdom), (1992).
